# Acute ClC-2 inhibition depolarizes E_GABA_ and increases neuronal intracellular chloride

**DOI:** 10.64898/2026.09.24.753897

**Authors:** Fu Hung Shiu, Volodymyr I. Dzhala, Kyle P. Lillis, Kevin J. Staley

**Affiliations:** Department of Neurology, Harvard Medical School and Massachusetts General Hospital, Boston, MA 02114

## Abstract

Neuronal intracellular chloride concentration shapes the polarity and strength of GABA_A_ receptor-mediated inhibition. ClC-2 is a hyperpolarization-activated chloride channel proposed to support chloride extrusion, but its acute contribution to E_GABA_ and neuronal chloride homeostasis remains incompletely understood. Here, we used the selective ClC-2 inhibitor AK-42 to test how acute ClC-2 inhibition affects GABAergic chloride signaling in organotypic hippocampal slice cultures. We combined targeted single-interneuron optogenetic activation of stCoChR-expressing GABAergic interneurons with gramicidin-perforated patch recordings to measure optogenetically evoked inhibitory postsynaptic currents and calculate E_GABA_ and we used SuperClomeleon imaging to measure neuronal intracellular chloride. AK-42 depolarized E_GABA_, with a larger effect when stimulation was delivered 2 s rather than 100 ms after voltage-step onset. This delay-dependent effect is consistent with greater engagement of voltage- and time-dependent ClC-2 activity during prolonged voltage steps. AK-42 also abolished the more hyperpolarized E_GABA_ normally measured at the 2-s stimulation delay and shifted baseline holding currents at hyperpolarized command potentials without measurably changing estimated resting membrane potential. SuperClomeleon imaging showed that AK-42 increased neuronal intracellular chloride in the presence of TTX, supporting a role for ClC-2 in basal chloride regulation. Together, these findings show that acute ClC-2 inhibition depolarizes E_GABA_, increases neuronal intracellular chloride, and alters the temporal dynamics of GABAergic chloride signaling, identifying ClC-2 as an acute regulator of synaptic GABAergic chloride signaling.

## Introduction

Fast synaptic inhibition in the brain is primarily mediated by GABA_A_ receptors, ligand-gated anion channels that conduct mainly chloride (Cl^-^) with small contribution of bicarbonate (HCO_3_−) under physiological conditions (Bormann, Hamill et al. 1987). The polarity and strength of GABAergic signaling depend not only on GABA_A_ receptor conductance, but also on the electrochemical driving force for Cl.

Even modest increases in intracellular CI^-^ ([Cl^−^]_i_) concentration can shift the GABA reversal potential (E_GABA_) toward more depolarized values. In the mature brain, E_GABA_ is typically slightly more negative than the resting membrane potential (RMP), resulting in hyperpolarizing or shunting inhibition (Staley and Mody 1992, Rivera, Voipio et al. 1999). During pathological activity, including seizures, impaired Cl^−^ extrusion or excessive Cl^−^ influx can transiently depolarize E_GABA_, weakening inhibition or converting GABAergic responses from inhibitory to depolarizing (Staley, Soldo et al. 1995, Lillis, Kramer et al. 2012, Sulis Sato, Artoni et al. 2017, Burman, Selfe et al. 2019). Therefore, understanding how neuronal Cl^−^ gradients are established and maintained is important for defining the mechanisms that preserve effective GABAergic inhibition.

Intra and extracellular [Cl^−^]_i_ concentrations are determined largely by Donnan exclusion (Glykys, Dzhala et al. 2014, Rahmati, Normoyle et al. 2021, Normoyle, Lillis et al. 2025). However, transmembrane ion flux alters local ion concentrations, so that neurons have the choice of changing volume or returning ion concentrations back to their steady state values (Staley 2024). This restoration of intracellular chloride concentration is commonly attributed to the coordinated actions of cation–chloride cotransporters, particularly the Cl^-^ extruding transporter KCC2 and the Cl-importing transporter NKCC1 (Kahle, Staley et al. 2008, Blaesse, Airaksinen et al. 2009). However, passive Cl^-^ conductances may also contribute to chloride regulation by permitting Cl^-^ movement according to its electrochemical gradient. Among the candidate channels involved in this process, ClC-2 is particularly relevant because it is activated at hyperpolarized membrane potential and may facilitate Cl^−^ movement when intracellular Cl^−^ is elevated such that the chloride reversal potential is positive to membrane potential (Staley 1994, Rinke, Artmann et al. 2010).

ClC-2 is an inwardly rectifying Cl channel widely expressed across tissues, with prominent expression in the brain and epithelia (Thiemann, Grunder et al. 1992, Cid, Montrose-Rafizadeh et al. 1995, Jentsch and Pusch 2018). In the brain, it is expressed in both neurons and glia (Smith, Clayton et al. 1995, Makara, Rappert et al. 2003), and one of its earliest proposed neuronal functions is to assist Cl extrusion after high chloride load (Staley 1994, Smith, Clayton et al. 1995, Staley, Smith et al. 1996). In ClC-2 knockout mice, pyramidal neurons and subsets of interneurons showed increased input resistance, hence excitability (Rinke, Artmann et al. 2010), suggesting ClC-2 is active at membrane potential. Furthermore, after a large, experimentally induced Cl^−^ load, neurons lacking ClC-2 recover more slowly, supporting a role for ClC-2 in accelerating Cl^−^ extrusion under conditions of elevated [Cl^−^]_i_ (Rinke, Artmann et al. 2010). In hippocampal circuits, ClC-2 has also been implicated in input-specific chloride regulation at parvalbumin-positive (PV) basket-cell synapses onto hippocampal pyramidal neurons, where it may help limit postsynaptic chloride accumulation during repetitive GABAergic transmission (Foldy, Lee et al. 2010). However, the contribution of ClC-2 to neuronal chloride homeostasis remains debated (Jentsch and Pusch 2018). Using computational modeling, Ratté and Prescott proposed that under physiological resting conditions ClC-2 primarily mediates Cl^−^ influx rather than extrusion and reduces neuronal excitability through its contribution to membrane conductance, with relatively little effect on steady-state intracellular Cl^−^ concentration (Ratte and Prescott 2011).

Defining the physiological function of ClC-2 has also been complicated by the limitations of constitutive knockout models. Although Clcn2-knockout mice have provided important information, developmental compensation, altered gene expression, and homeostatic plasticity may reduce or obscure the immediate consequences of ClC-2 loss. These compensatory mechanisms may partly explain why Clcn2-knockout mice exhibit relatively subtle neuronal phenotypes and lack a robust spontaneous epilepsy phenotype (Bosl, Stein et al. 2001, Nehrke, Arreola et al. 2002, Blanz, Schweizer et al. 2007). Similarly, although several *CLCN2* variants have been identified in patients with epilepsy, human genetic studies have not established a consistent causal relationship between *CLCN2* loss of function and epilepsy syndromes (Planells-Cases and Jentsch 2009, Niemeyer, Cid et al. 2010). Consequently, the immediate contribution of ClC-2 to neuronal chloride homeostasis and GABAergic signaling remains incompletely understood.

Together, these studies suggest that ClC-2 can contribute to neuronal chloride regulation, but they do not establish whether ongoing ClC-2 activity acutely maintains E_GABA_ and intracellular chloride. This question is difficult to resolve with constitutive knockout models, because developmental compensation may mask the immediate effects of ClC-2 loss. It is also difficult to infer the contribution of ClC-2 to neuronal chloride homeostasis from ClC-2 current measurements alone, because ClC-2-mediated chloride movement depends on membrane voltage, and the ratio of intra-to extracellular Cl^−^-i.e. the electrochemical driving force for Cl^−^. Direct measurements of E_GABA_ and neuronal [Cl^−^]_i_ are therefore needed to determine how acute ClC-2 inhibition affects GABAergic chloride signaling.

Acute pharmacological inhibition provides a direct way to test this question. Koster et al. recently developed AK-42, a potent and selective small-molecule inhibitor of ClC-2 that provides a pharmacological approach for addressing these questions (Koster, Reese et al. 2020). AK-42 inhibits ClC-2 with nanomolar potency and displays more than 1,000-fold selectivity over the closely related ClC-1 channel, while showing minimal activity against a broad panel of brain-expressed ion channels, receptors, and transporters. In hippocampal CA1 pyramidal neurons, AK-42 acutely and reversibly inhibited ClC-2-mediated currents, whereas no effect was observed in tissue from *Clcn2*-knockout mice. Subsequent cryo-EM studies visualized AK-42 bound to ClC-2 and showed that AK-42 occupies the extracellular entryway or outer vestibule of the chloride-permeation pathway, providing structural support for a direct pore-blocking mechanism (Ma, Wang et al. 2023, Xu, Neelands et al. 2024).

In this study, we used AK-42 to determine how acute ClC-2 inhibition affects neuronal chloride regulation and GABAergic signaling in organotypic hippocampal slice cultures. If ClC-2 normally helps limit neuronal Cl^−^ accumulation, then blocking ClC-2 should increase [Cl^−^]_i_ and shift E_GABA_ toward more depolarized values. To test this, we measured optogenetically evoked E_GABA_ using gramicidin-perforated patch recordings and measured neuronal [Cl^−^]_i_ using SuperClomeleon, a genetically encoded ratiometric fluorescent chloride indicator (Grimley, Li et al. 2013).

We also tested whether ClC-2 contributes to time-dependent changes in the measured E_GABA_ during voltage-step protocols. Because ClC-2 activates gradually after membrane hyperpolarization (Thiemann, Grunder et al. 1992), GABAergic currents were evoked either 100ms or 2s after voltage-step onset. This allowed us to determine whether prolonged voltage-step onset reveals a ClC-2-dependent component of the GABAergic current–voltage relationship.

Together, these electrophysiological and imaging approaches allowed us to determine that acute ClC-2 inhibition depolarizes E_GABA_, increases neuronal [Cl^−^]_i_, and abolishes a stimulation-delay-dependent hyperpolarization of E_GABA_.

## Experimental design and methods

All animal-use protocols were in accordance with the guidelines of the National Institutes of Health and the Massachusetts General Hospital Center for Comparative Medicine on the use of laboratory animals. All protocols were approved by the MGH Institutional Animal Care and Use Committee (IACUC).

### Organotypic hippocampal slice (OHSC) preparation

Postnatal day 7 of DLX-cre mice (for E_GABA_ optogenetic experiment) or WT mice (Cl imaging) were used to prepare organotypic hippocampal slice culture (OHSCs) using the rocking plate technique (Romijn, de Jong et al. 1988, Berdichevsky Y 2017). Isolated hippocampi were cut into 400µm slices on a McIlwain tissue chopper (Mickle Laboratory Engineering). Slices were transferred to coverslips, which were placed in glass-bottomed six-well plates (catalog #P06-1.5H-N, CellVis). Both culture configurations were incubated at 5% CO_2_, 37°C in Neurobasal-A growth medium supplemented with 2% B27, 500 μm GlutaMAX, and 0.03 mg/ml gentamycin (all from Thermo Fisher Scientific). The growth medium was changed every 3–4 d. Slices were used for experiments between day *in vitro* (DIV) 9 and DIV 16.

### Virus approach

The virus was delivered on the day of slicing. For E_GABA_ experiment, we utilized optogenetics to activate DLX-expressing interneurons with an excitatory opsin (StCoChR) that is capable of single-neuron activation (Forli, Pisoni et al. 2021, Printz, Patil et al. 2023). Viral solution AAV2/9-hEF1α-dlox-stCoChR-mScarlet (v934-1, titer, 2.4 × 10^13^ vg/mL) (2 µl/ml culture media) was added into the cell culture media on the day of slicing. The viral solution was constructed from the Viral Vector Facility of the Neuroscience Center Zurich, through a generous donation from Ofer Yizhar. For intracellular chloride imaging experiments, AAV2/9-Syn-SuperClomeleon (synthesized by the Gene Transfer Vector Core at MGH, 8.9 × 10^13^ vg/mL) was added to the culture medium at 2 µl/ml on the day of slicing (Costine-Bartell, Martinez-Ramirez et al. 2023).

### CLC-2 inhibitor, AK42

AK-42 (SML3933, MilliporeSigma) was dissolved and aliquoted in DMSO. Aliquots were stored at −20°C and used within 1 month. On the day of recording, AK-42 was diluted into aCSF to a final concentration of 2.5 µM.

### Patch recording

For gramicidin-perforated patch recordings, the internal solution contained 150 mM KCl and 10 mM HEPES, adjusted to pH 7.4 with KOH and 290–300 mOsm. Gramicidin was added freshly on the day of recording to a final concentration of 50 µg/mL, and the internal solution was thoroughly mixed before pipette filling (Raveendran, Pressey et al. 2025). Recordings were performed after gradual perforation, with access resistance allowed to stabilize below 50 MΩ. Cells were excluded if access resistance changed abruptly, indicating accidental membrane rupture and transition to the whole-cell configuration. At the end of each experiment, the membrane was intentionally ruptured to establish the whole-cell configuration, allowing the neuron to be filled with dye for subsequent examination of its morphology.

Whole-cell patch-clamp recordings were used to validate stCoChR. For these recordings, patch pipettes were filled with an internal solution containing the following, in mM: 135 potassium gluconate, 2 MgCl_2_, 0.1 CaCl_2_, 1 EGTA, 2 Na_2_ATP, and 10 HEPES. The solution was adjusted to pH 7.25 with KOH and 290–300 mOsm.

During recordings, kynurenic acid (KYNA; 3 mM), a broad-spectrum ionotropic glutamate receptor antagonist, was included in the bath solution to block glutamatergic synaptic transmission.

### E_GABA_ measurement

Postsynaptic neurons in OHSCs were voltage-clamped at a holding potential of -50 mV and stepped to command potentials from −100 to −50mV in 10mV increments. Steps were 6 seconds in duration. The interval between steps was 5 seconds. At each command potential, GABAergic postsynaptic currents were evoked by spatially targeted optogenetic stimulation of individual presynaptic interneurons expressing st-CoChR. St-CoChR was activated using laser targeted path scanning via a 920 nm bandpass filter. These optical stimuli were of ≤5 ms duration and were delivered 100ms or 2s after the onset of the voltage step. For each command potential, the peak optogenetically evoked current amplitude was measured relative to the pre-stimulus baseline. Current–voltage relationships were generated by plotting the evoked current amplitude against the corresponding command potential. E_GABA_ was estimated from these current–voltage relationships using the Goldman– Hodgkin–Katz constant-field equation (Hodgkin and Katz 1949), as described by Bormann et al (Bormann, Hamill et al. 1987).

### Chloride imaging

Intracellular chloride was assessed using SuperClomeleon (sCLM) (Grimley, Li et al. 2013), a genetically encoded ratiometric chloride indicator derived from Clomeleon (Kuner and Augustine 2000). sCLM contains Cerulean, a chloride-insensitive cyan fluorescent protein (CFP) and a chloride-sensitive Topaz-derived yellow fluorescent protein (YFP) joined by a short linker, allowing fluorescence resonance energy transfer (FRET)-based measurement of intracellular chloride. Because chloride quenches YFP fluorescence, decreases in the YFP/CFP fluorescence ratio were interpreted as increases in intracellular chloride.

For chloride imaging experiments, OHSCs expressing sCLM were transferred to a submerged imaging chamber and continuously perfused with oxygenated ACSF at 34°C. TTX (1069, Tocris Bioscience, 1µM) was added to prevent any neuronal activities that might cause activity-dependent Cl_i_ changes. Two-photon imaging was performed on an Olympus Fluoview 1000 MPE microscope. Excitation of 860nm was used in a mode-locked titanium-sapphire laser (MaiTai, Spectra Physics. Emitted light passed through a dichroic mirror and was bandpass filtered through 480 ± 15 nm (D480/30) for CFP and a 535 ± 20-nm filter (D535/40) for YFP (FV10-MRCYR/XR). Imaging settings, including laser power, detector gain, pixel dwell time, and scan resolution, were kept constant across experimental conditions within each experiment.

Regions of interest were drawn around neuronal somata and, when clearly resolved, proximal neurites. For experiments involving AK-42, baseline sCLM images were acquired before drug application, followed by imaging after bath application of AK-42 for under 30 minutes under identical acquisition settings. For the control group, baseline sCLM images were acquired using the same acquisition settings, followed by a second imaging session 30 min later (Baseline + 30 min).

The absolute intracellular chloride concentration was calculated as follows: The YFP/CFP ratio (R) was converted to intracellular chloride concentration using the calibration relationship: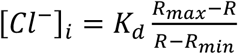, where (R) is the measured YFP/CFP ratio, (R_max_) is the ratio obtained under chloride-free conditions, (R_min_) is the ratio obtained under saturating chloride conditions, and (K_d_) is the chloride dissociation constant for sCLM (Grimley, Li et al. 2013, Rahmati, Normoyle et al. 2021, Bahari, Dzhala et al. 2024).

### Statistical analysis

Statistical analyses were performed using GraphPad Prism. For comparisons of E_GABA_ before and after AK-42 application, paired t tests were used. When comparisons of EGABA and ΔE_GABA_ were made at both 100-ms and 2-s stimulation delays, p values were corrected using the Holm-Šídák method.

Optogenetically evoked GABAergic current amplitudes across voltage steps were analyzed using two-way repeated-measures ANOVA, Baseline-versus-treatment comparisons at individual voltage steps were performed using Šídák corrections for multiple comparisons.

Changes in resting membrane potential were calculated using the same ΔE_GABA_ approach and compared between stimulation delays or treatment conditions as indicated. Data are reported as mean ± SD unless otherwise stated. Statistical significance was defined as p ≤ 0.05.

## Results

### Single-interneuron optogenetic stimulation with stCoChR reliably evokes postsynaptic GABAergic currents

To measure GABAergic responses evoked by individual interneurons in OHSCs, we used a cre-dependent viral strategy to selectively express the excitatory opsin stCoChR (Forli, Pisoni et al. 2021, Printz, Patil et al. 2023), together with an mScarlet reporter, in DLX-lineage interneurons of DLX-Cre mice. This approach allowed presynaptic interneurons to be optically stimulated while recordings were obtained from nearby postsynaptic visually identified pyramidal cells. Compared with bulk electrical or field stimulation, targeted optical stimulation provides greater control over the recruited inhibitory input and reduces the simultaneous activation of multiple GABAergic fibers.

Following viral transduction for a week, mScarlet-positive interneurons were readily identified in OHSCs and targeted for whole-cell patch-clamp recording and optical stimulation (**Figure 1A**). We first determined whether brief two-photon stimulation was sufficient to activate stCoChR-expressing interneurons. During current-clamp recordings, stCoChR-expressing interneurons were filled with Cascade Blue™ to confirm the identity and location of the recorded cell **(Figure 1B–C**). Two-photon stimulation at 920 nm using 5-ms pulses delivered at 0.33 Hz reliably evoked action potentials in the recorded interneuron (n=3 neurons) (**Figure 1D**) as described previously (Forli, Pisoni et al. 2021).

**Figure 1.**
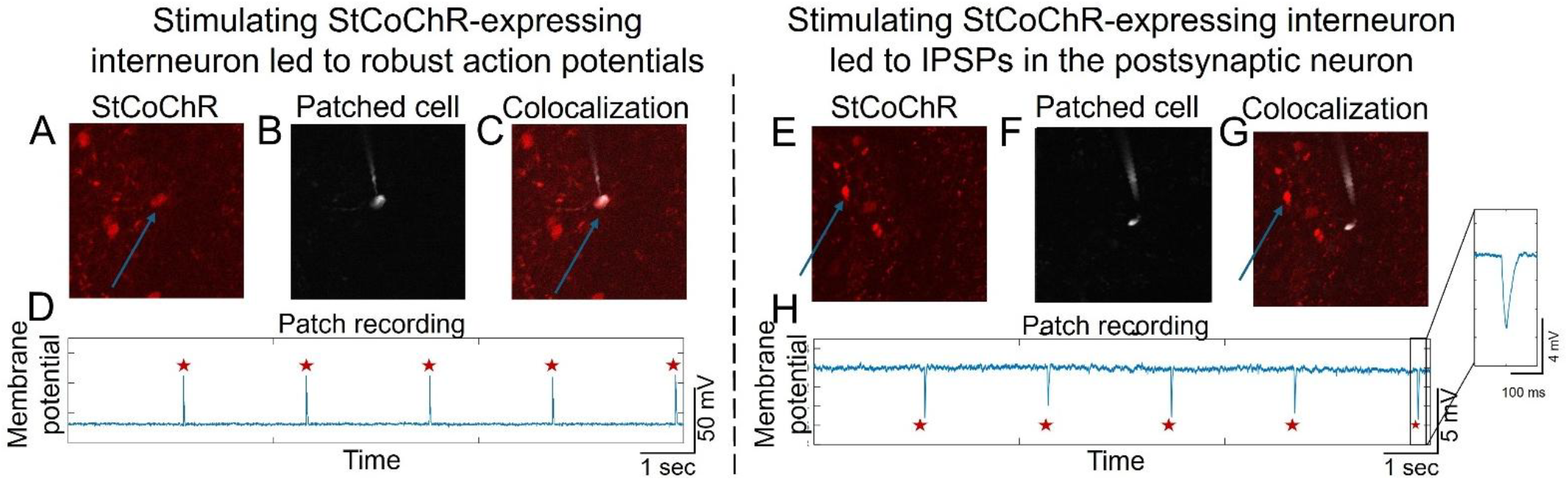
Optical stimulation of stCoChR-expressing interneurons evokes action potentials and inhibitory synaptic responses in postsynaptic neurons. **(A–C)** Representative stCoChR-expressing interneuron filled with Cascade Blue™ during whole-cell recording. **(D)** Two-photon stimulation at 920 nm (0.33 Hz, 5-ms pulses) reliably evoked action potentials. **(E–G)** Representative Cascade Blue™-filled postsynaptic neuron adjacent to a stCoChR-expressing interneuron. **(H)** Optical stimulation of the presynaptic interneuron evoked time-locked inhibitory postsynaptic potentials. Red stars indicate optical stimulation. The inset shows a magnified view of a representative IPSP.

We next examined whether stimulation of a single stCoChR-expressing interneuron could evoke synaptic responses in a neighboring neuron. Postsynaptic neurons located near an identified stCoChR-positive interneuron were targeted for whole-cell current-clamp recording and filled with Cascade Blue™ (**Figure 1E–G**). Optical stimulation of the presynaptic interneuron evoked time-locked inhibitory postsynaptic potentials in the neighboring postsynaptic neuron (n= 3 neurons) (**Figure 1H**). These responses demonstrate that this approach can reliably activate individual interneurons and measure the corresponding postsynaptic GABAergic responses in neighboring visually-identified pyramidal cells.

### ClC-2 inhibition with AK-42 depolarizes optogenetically evoked E_GABA_ and alters its dependence on stimulation delay

We next examined whether acute pharmacological inhibition of ClC-2 depolarizes the reversal potential of optogenetically evoked GABAergic responses. E_GABA_ was measured using gramicidin-perforated patch recordings to preserve the endogenous intracellular chloride gradient. During each voltage-step protocol, a presynaptic stCoChR-expressing interneuron was optically stimulated either 100-ms or 2-s after voltage-step onset. Representative traces are shown in **Figure 2A**.

**Figure 2.**
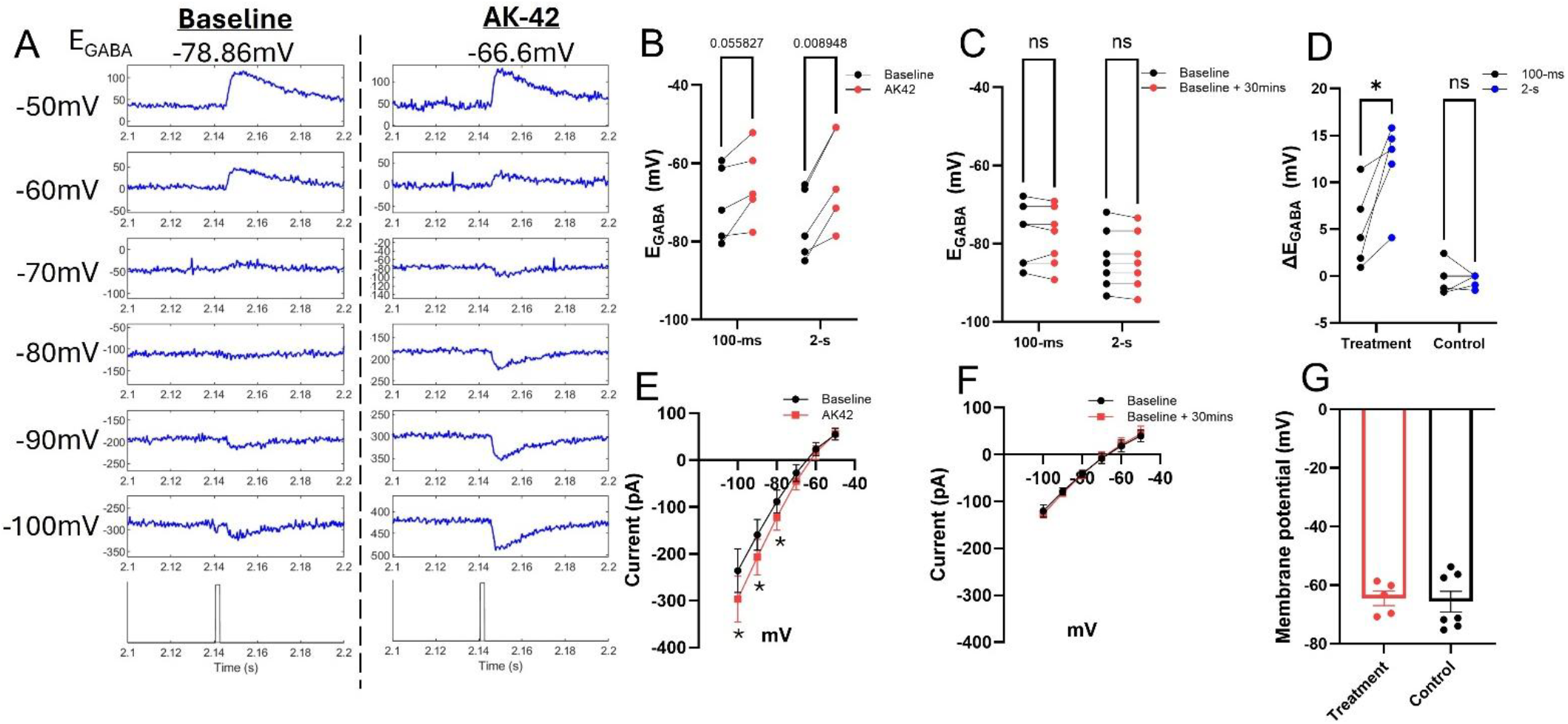
Inhibition of ClC-2 with AK-42 depolarizes optogenetically evoked E_GABA_. **(A)** Representative current–voltage relationships showing an AK-42-induced shift in E_GABA_ from −78.6 to −66.6 mV. Representative traces were low-pass filtered offline using a Gaussian filter with a −3 dB cutoff frequency of 1 kHz. **(B)** Paired measurements showing depolarization of E_GABA_ after AK-42 application when optical stimulation was delivered 100-ms (left) or 2-s (right) after voltage-step onset. **(C)** E_GABA_ remained stable in time-matched control recordings without AK-42. **(D)** ΔE_GABA_, calculated as the post-treatment value minus baseline, was significantly greater at the 2-s delay than at the 100-ms delay after AK-42 treatment (left), whereas no delay-dependent difference was observed in time-matched control recording(right). **(E)** Baseline holding-current I–V relationships measured before and after AK-42 application. AK-42 shifted holding currents in the negative direction at hyperpolarized command potentials, with significant differences at −100, −90, and −80 mV. **(F)** Baseline holding-current I–V relationships remained similar in time-matched control recording. **(G)** Estimated resting membrane potential calculated from the holding-current I– V relationship did not differ significantly between AK-42-treated cells and time matched controls.

Baseline E_GABA_ was first measured in aCSF at both stimulation delays. AK-42 (2.5µM) was then bath-applied for 30 min, and the measurements were repeated. When optical stimulation was delivered 100 ms after voltage-step onset, AK-42 shifted E_GABA_ from −70.31 ± 9.74mV at baseline to −65.23 ± 9.78mV after AK-42, corresponding to a 5.09 ± 4.26mV depolarizing shift that approached significance (n = 5 paired recordings, paired t test; t(4) = 2.67, adjusted p = 0.0558; **Figure 2B, left**). When optical stimulation was delivered 2-s after voltage-step onset, AK-42 significantly depolarized E_GABA_ from −75.65 ± 9.08mV at baseline to −63.65 ± 12.47mV after AK-42, corresponding to a 12 ± 4.65mV depolarizing shift (n = 5 paired recordings, paired t test with Holm–Šídák correction; t(4) = 5.77, adjusted p = 0.0090; **Figure 2B, right**).

To determine whether these E_GABA_ shifts resulted from AK-42 rather than recording duration or repeated voltage-step protocols, we performed time-matched control recordings in aCSF without drug application. E_GABA_ remained stable between baseline and baseline + 30 mins at both stimulation delays (n = 7 paired recordings, 100-ms: adjusted p = 0.57; 2-s: adjusted p = 0.34; **Figure 2C**).

We next calculated ΔE_GABA_ as the E_GABA_ from post-treatment (either E_GABA_ obtained from optical stimulation 100-ms or 2-s after voltage step) minus the corresponding baseline value. In AK-42-treated cells, ΔE_GABA_ was significantly greater when optical stimulation was delivered 2-s after voltage-step onset than when stimulation was delivered after 100-ms. AK-42 depolarized E_GABA_ by 5.09 ± 4.26mV at the 100-ms delay and by 12 ± 4.65mV at the 2-s delay, corresponding to a significantly larger depolarizing shift at the longer stimulation delay (n= 5 paired recordings, paired t test with Holm-Šídák correction; t(4) = 3.57, adjusted p = 0.0464; **Figure 2D left**). No delay-dependent difference in ΔE_GABA_ was observed in time-matched control recordings (100-ms: −0.33 ± 1.45 mV; 2-s: −0.35 ± 0.62 mV; n = 7 paired recordings; paired t test with Holm–Šídák correction; t(6) = 0.043, adjusted p = 0.9670; **Figure 2D right**).

Because ClC-2 is voltage dependent, we also examined whether AK-42 changed the baseline holding-current I–V relationship during the voltage-step protocol. AK-42 shifted holding currents in the negative direction at hyperpolarized command potentials (two-way repeated-measures ANOVA; F(5,20) = 4.60, p = 0.0059; **Figure 2E**). Post hoc comparisons showed significantly more negative holding currents after AK-42 at −100, −90, and −80 mV, but not at more depolarized command potentials. This voltage-dependent change is consistent with inhibition of a hyperpolarization-activated ClC-2 conductance. In contrast, time-matched control recordings showed stable holding-current I–V relationships over the same recording interval (**Figure 2F**).

Despite this voltage-dependent change in holding current, AK-42 did not significantly change the estimated resting membrane potential (RMP) calculated from the 0-current condition in the command potential I–V relationship. The AK-42-induced change in estimated RMP was also not significantly different from the change observed in time-matched controls (**Figure 2G**). Thus, AK-42 altered baseline holding current at hyperpolarized voltages without producing a detectable shift in estimated RMP. These data show AK-42 depolarizes E_GABA_ through effects on chloride regulation rather than through a nonspecific shift in RMP.

### AK-42 shifts optogenetically evoked GABAergic currents

We also analyzed the optogenetically-evoked GABAergic current amplitudes used to generate the current–voltage relationships. For 100-ms stimulation (**Figure 3A**), AK-42 produced a significant main effect on current amplitude across voltage steps (F(1,4) = 16.61, p = 0.015). Baseline-versus-AK-42 differences were significant at −100 mV and −90 mV, but not at more depolarized command potentials (Šídák-corrected comparisons; −100 mV: p = 0.0050; −90 mV: p = 0.047). Time-matched control recordings showed no significant changes (**Figure 3B**). This increased amplitude of the IPSC at negative command potentials supports the idea that ClC2 is depleting intracellular chloride at negative command potentials, resulting in an E_GABA_ that is closer to the command potential. This reduces the driving force for the IPSC in control conditions. Blocking ClC2 with AK42 prevents this decrement in the driving force for the IPSC.

**Figure 3.**
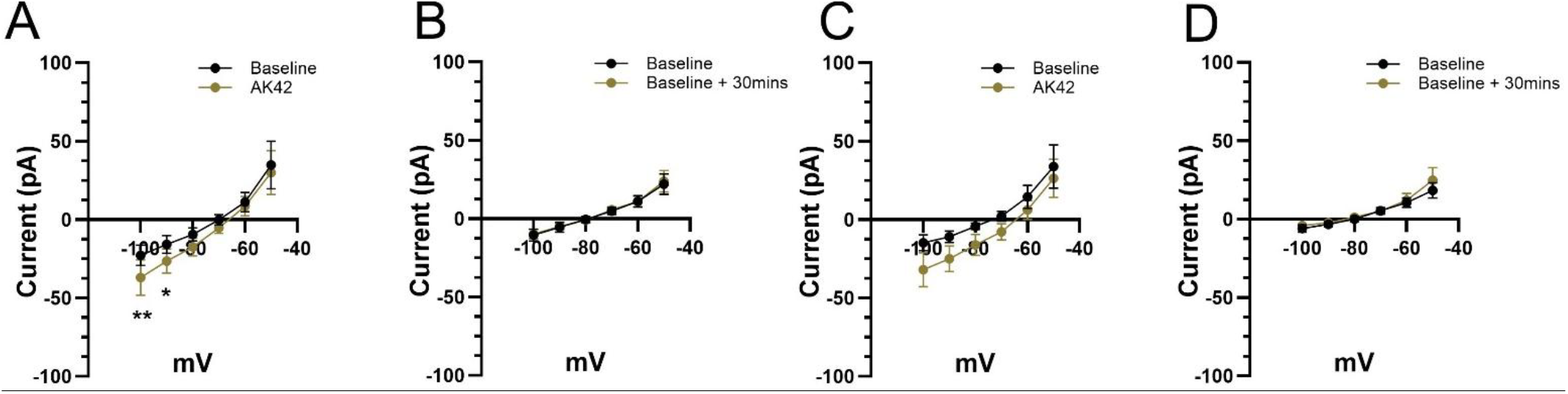
AK-42 shifts optogenetically evoked GABAergic current amplitudes. (A) Current–voltage relationships measured before and after AK-42 application with stimulation delivered 100 ms after voltage-step onset. AK-42 increased the magnitude of negative currents at hyperpolarized command potentials, with significant differences at −100 and −90 mV. (B) Time-matched control recordings at the 100-ms delay showed no significant change. (C) Current–voltage relationships measured before and after AK-42 application with stimulation delivered 2 s after voltage-step onset. AK-42 shifted current amplitudes in the negative direction, with a trend toward significance. (D) Time-matched control recordings at the 2-s delay showed no significant change.

For 2-s stimulation (**Figure 3C**), AK-42 shifted current amplitudes in the negative direction and showed a trend toward a treatment effect across voltage steps (F(1,4) = 5.94, p = 0.0714). Time-matched control recordings showed no significant changes when optical stimulation was delivered 2-s after voltage-step onset (**Figure 3D**).

### Optical stimulation at 2-s delay produces a more hyperpolarized E_GABA_, and AK-42 abolishes this difference

To determine why the AK-42-induced ΔE_GABA_ was greater at the 2-s stimulation delay, we directly compared the absolute E_GABA_ values measured at 100-ms and 2-s. Under baseline conditions, E_GABA_ was significantly more hyperpolarized when stimulation was delivered after 2 s after voltage-step onset than when stimulation was delivered after 100 ms. Baseline E_GABA_ was −70.31 ± 9.74mV at the 100-ms delay and −75.65 ± 9.08mV at the 2-s delay, corresponding to a 5.34 ± 1.52mV hyperpolarization at the longer delay (n = 5 paired recordings, paired t test with Holm-Šídák correction; t(4) = 7.86, adjusted p = 0.0028; **Figure 4A left**). Following AK-42 application, this delay-dependent difference was no longer present. E_GABA_ was −65.23 mV at the 100-ms delay and −63.65 mV at the 2-s delay after AK-42 treatment (n = 5 paired recordings, adjusted p = 0.45; **Figure 4A right**).

**Figure 4.**
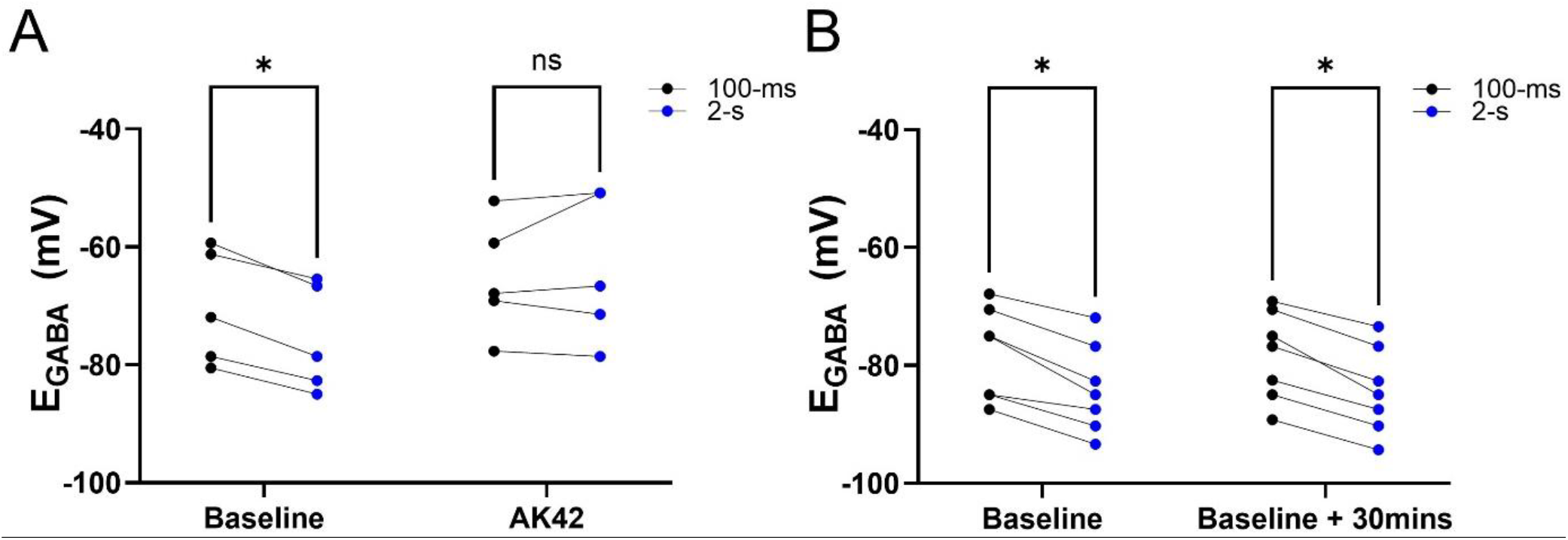
Optical stimulation at 2-s yields a more hyperpolarized EGABA than stimulation at 100-ms, and AK-42 abolishes this difference. (A) At baseline, E_GABA_ was significantly more hyperpolarized when optical stimulation was delivered 2-s after voltage-step onset than when delivered after 100-ms. This difference was no longer present following AK-42 application. (B) In time-matched aCSF controls, EGABA remained more hyperpolarized at the 2-s delay during both baseline and baseline + 30mins measurements.

In time-matched control recordings, E_GABA_ remained more hyperpolarized at the 2-s delay than at the 100-ms delay during both baseline and baseline + 30mins measurement. This delay-dependent difference was significant at baseline (n = 7 paired recordings; paired t test with Holm–Šídák correction; t(6) = 6.58, adjusted p = 0.00059; **Figure 4B**, left) and remained significant after the 30 mins interval (n = 7 paired recordings; paired t test with Holm–Šídák correction; t(6) = 8.50, adjusted p = 0.00029; **Figure 4B**, right).

These results demonstrate that a longer stimulation delay produces a more hyperpolarized E_GABA_ measurement under control conditions. In contrast, AK-42 removed this delay-dependent difference.

### AK-42 increases intracellular chloride measured with sCLM imaging

To test whether AK-42-induced E_GABA_ depolarization was accompanied by increased neuronal [Cl^−^]_i_, we measured neuronal [Cl^−^]_i_ using sCLM. Representative sCLM images were acquired before and after AK-42 application or during time-matched control group (**Figure 5A**).

**Figure 5.**
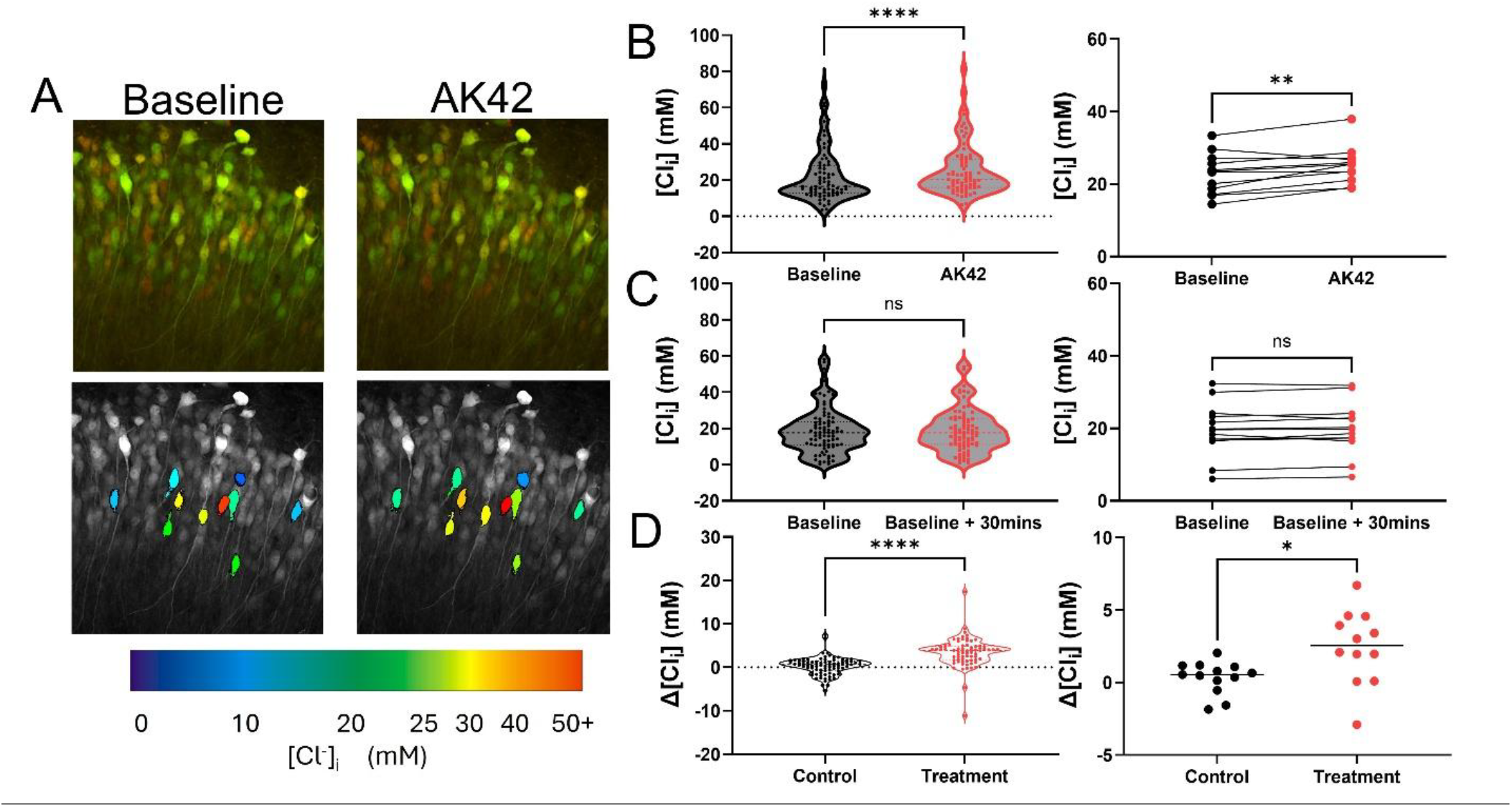
AK-42 increases [Cl^−^]_i_ measured with sCLM. (A) Representative sCLM images before and after AK-42 application (Top). The same fields of view with false-color [Cl^−^]_i_ maps, with blue indicating lower [Cl^−^]_i_ and red indicating higher [Cl^−^]_I_ (bottom). AK-42 shifted the false-color signal toward higher [Cl^−^]_i._ (B) Estimated neuronal [Cl^−^]_i_ before and after AK-42 application, shown for individual neurons (left) and slice averages (right). AK-42 significantly increased estimated [Cl^−^]_i_ at both the neuron and slice levels. (C) Estimated neuronal [Cl^−^]_i_ during time-matched control recordings, shown for individual neurons (left) and slice averages (right). Control recordings showed no significant change in estimated [Cl^−^]_i_. (D) Δ[Cl^−^]_i_, calculated as the AK42 or baseline + 30mins value minus baseline, in AK-42-treatment and control groups. Individual-neuron Δ[Cl^−^]_i_ is shown on the left, and slice-level Δ[Cl^−^]_i_ is shown on the right. AK-42 produced a larger increase in Δ[Cl^−^]_i_ than controls at both levels.

AK-42 increased estimated neuronal [Cl^−^]_i_ after drug application (**Figure 5B**). At the individual-neuron level, estimated [Cl^−^]_i_ increased from 22.03 ± 14.39mM at baseline to 25.53 ± 14.38mM after AK-42 treatment (n = 73 neurons; paired t test, t(72) = 9.04, p < 0.0001; **Figure 5B, left**). This increase remained significant when values were averaged by slice, with estimated [Cl^−^]_i_ increasing from 22.80 ± 5.62mM at baseline to 25.26 ± 5.062mM after AK-42 treatment (n = 12 slices; paired t test, t(11) = 3.35, p = 0.0065; **Figure 5B, right**).

In contrast, time-matched control group showed no significant increase in estimated [Cl^−^]_i_ (**Figure 5C**). At the individual-neuron level, estimated [Cl^−^]_i_ was 18.75 ± 12.52mM at baseline and 19.07 ± 12.11mM at baseline + 30mins (n = 83 neurons; paired t test, t(82) = 1.44, p = 0.154; **Figure 5C, left**). At the slice level, estimated [Cl^−^]_i_ was 19.52 ± 7.34mM at baseline and 19.87 ± 7.16mM at baseline + 30mins (n = 13 slices; paired t test, t(12) = 1.13, p = 0.281; **Figure 5C, right**).

We next quantified Δ[Cl^−^]_i_ as the AK42 or baseline + 30mins minus baseline. Treatment group produced a larger increase in Δ[Cl^−^]_i_ than control group at the individual-neuron level (Treatment: 3.50 ± 0.39 mM, n = 73 neurons; Control: 0.32 ± 0.22 mM, n = 83 neurons; Welch’s t test, t(116.3) = 7.12, p < 0.0001; **Figure 5D**). This AK-42-associated increase was also significant at the slice level (Treatment: 2.46 ± 0.73 mM, n = 12 slices; Control: 0.34 ± 0.30 mM, n = 13 slices; Welch’s t test, t(14.7) = 2.66, p = 0.0181; **Figure 5E**).

## Discussion

In this study, we combined single neuron optogenetic stimulation of GABAergic interneurons, gramicidin-perforated patch recording, and chloride imaging to examine the acute contribution of ClC-2 to GABAergic chloride signaling in OHSCs. Acute ClC-2 inhibition with AK-42 depolarized optogenetically evoked E_GABA_, abolished the stimulation-delay-dependent hyperpolarization of E_GABA_, and increased neuronal [Cl^−^]_i_. Together, these findings support a role for ClC-2 in limiting [Cl^−^]_i_ accumulation and maintaining hyperpolarized GABAergic signaling.

Previous studies have implicated ClC-2 in neuronal chloride regulation, particularly after intracellular chloride loading or during specific GABAergic inputs. ClC-2 expression was shown to shift E_Cl_ in a hyperpolarizing direction (Staley, Smith et al. 1996), and neurons from Clcn2-knockout mice show longer recovery after experimentally induced chloride loading (Rinke, Artmann et al. 2010). However, most prior work inferred the role of ClC-2 from ClC-2-dependent currents, GABA_A_ current properties, or recovery after imposed chloride loads.

Here, using the selective ClC-2 inhibitor AK-42 (Koster, Reese et al. 2020), we show that acute ClC-2 inhibition is sufficient to depolarize optogenetically evoked E_GABA_ and increase neuronal [Cl^−^]_i_ in OHSCs. Because E_GABA_ measurement and Cl imaging were performed in the presence of KYNA or TTX respectively, the AK-42 induced depolarization of E_GABA_ and increase of [Cl^−^]_i_ does not require ongoing network activity, supporting a role for ClC-2 in basal chloride regulation in OHSC.

A second major finding is that E_GABA_ depended on the timing of optogenetic stimulation during the voltage-step protocol. Under baseline conditions, stimulation delivered 2 s after voltage-step onset produced a more hyperpolarized E_GABA_ than stimulation delivered after 100 ms, and this delay-dependent difference persisted in time-matched controls. AK-42 eliminated this difference, suggesting that the more hyperpolarized E_GABA_ measured after prolonged voltage-step onset depends on ClC-2 activity. Consistent with these E_GABA_ changes, AK-42 also increased optogenetically evoked GABAergic current amplitudes in the negative direction, with a significant effect during the 100-ms protocol and a trend during the 2-s protocol. Although previous work showed that ClC-2 can regulate GABA_A_ current rectification and time-dependent IPSC behavior especially in PV basket cell inputs (Foldy, Lee et al. 2010), our results extend those findings by showing that ClC-2 can also shape the apparent E_GABA_.

This timing dependence is consistent with the voltage- and time-dependent gating of ClC-2. During the 2-s delay, the postsynaptic neuron is held at the command potential long enough for slower voltage-dependent chloride conductances to influence the GABAergic current– voltage relationship. In contrast, stimulation after 100-ms likely samples the GABAergic response before this delayed ClC-2-dependent component is fully activated. These findings also have methodological implications: E_GABA_ estimates should not always be treated as fixed values independent of the voltage-step protocol. Stimulation timing should be kept constant across experimental groups, particularly when comparing conditions that may affect ClC-2 activity, intracellular chloride, or membrane conductance.

The AK-42 change in baseline holding current provides additional evidence that ClC-2 contributes during the voltage-step protocol. AK-42 shifted holding currents in the negative direction at hyperpolarized command potentials, consistent with inhibition of a hyperpolarization-activated ClC-2 conductance. However, this effect was not accompanied by a detectable shift in estimated RMP, indicating that the E_GABA_ depolarization cannot be explained by a change of RMP. Instead, the holding-current result supports the idea that ClC-2 contributes to voltage-dependent membrane conductance while also influencing neuronal chloride homeostasis.

Future studies can extend these findings by testing whether acute ClC-2 inhibition similarly depolarizes E_GABA_ and increases neuronal [Cl^−^]_i_ in vivo. It will also be important to define whether ClC-2 contributes to chloride regulation in a region- and cell type-specific manner, because ClC-2 has been strongly implicated in CA1/CA3 pyramidal neurons (Smith, Clayton et al. 1995) and PV basket cell inputs (Foldy, Lee et al. 2010), whereas its functional contribution across other hippocampal regions and different interneuron populations remains incompletely defined. Combining targeted stCoChR activation with interneuron subtype-specific strategies and chloride imaging could help determine whether ClC-2 regulates GABAergic signaling in an input-dependent manner.

In summary, acute inhibition of ClC-2 with AK-42 depolarized optogenetically evoked E_GABA_, increased neuronal [Cl^−^]_i_, and abolished a stimulation-delay-dependent hyperpolarization of E_GABA_. These findings demonstrate ClC-2 contributes to chloride homeostasis in OHSC neurons and helps maintain hyperpolarized GABAergic signaling. They also show that ClC-2 can shape E_GABA_ estimates during voltage-step protocols, emphasizing the need to consider voltage and time-dependent chloride conductance when interpreting GABAergic reversal potentials.

